# OptIC Notch reveals mechanism that regulates receptor interactions with CSL

**DOI:** 10.1101/2023.02.20.528063

**Authors:** Jonathan M. Townson, Maria J. Gomez-Lamarca, Carmen Santa Cruz Mateos, Sarah J. Bray

## Abstract

Active Notch signalling is elicited through receptor-ligand interactions that result in release of the Notch intracellular domain (NICD), which translocates into the nucleus. NICD activates transcription at target genes forming a complex with the DNA-binding transcription factor CSL (CBF1/Su(H)/Lag-1) and co-activator Mastermind. Despite this, CSL lacks its own nuclear localisation sequence, and it remains unclear where the tripartite complex is formed. To probe mechanisms involved, we designed an optogenetic approach to control NICD release (OptIC-Notch) and monitored consequences on complex formation and target gene activation. Strikingly we observed that, when uncleaved, OptIC-Notch sequestered CSL in the cytoplasm. Hypothesising that exposure of a juxta membrane ΦWΦP motif is key to sequestration, we masked this motif with a second light sensitive domain in OptIC-Notch{ω}, which was sufficient to prevent CSL sequestration. Furthermore, NICD produced by light-induced cleavage of OptIC-Notch or OptIC-Notch{ω} chaperoned CSL into the nucleus and induced target gene expression, showing efficient light controlled activation. Our results demonstrate that exposure of the ΦWΦP motif leads to CSL recruitment and suggest this can occur in the cytoplasm prior to nuclear entry.

**Summary statement:** Light controlled exposure of the Notch ΦWΦP motif leads to CSL sequestration and suggest this interaction could occur prior to nuclear entry during normal signalling.

## Introduction

The DNA binding protein CSL (an acronym for CBF-1/RBPJ-κ in mammals, Suppressor of Hairless in *Drosophila melanogaster*, Lag-1 in *Caenorhabditis elegans*) is the core transcription factor in the Notch pathway (Bray, 2006; Kopan and Ilagan, 2009; Kovall and Blacklow, 2010). When Notch interacts with ligands on neighbouring cells, a conformational change triggers successive cleavages that release the Notch intracellular domain (NICD), which contains several nuclear localization signals and rapidly enters the nucleus (Sprinzak and Blacklow, 2021). NICD forms a complex with CSL, an interaction that creates a binding groove for the co-activator Mastermind (Mam) (Choi et al., 2012; Nam et al., 2006; Wilson and Kovall, 2006). As well as its’ role in this tripartite activation complex, CSL is also part of several co-repressor complexes (Collins et al., 2014; Maier et al., 2011; Oswald and Kovall, 2018; Tabaja et al., 2017; Vanderwielen et al., 2011; Wacker et al., 2011; Yuan et al., 2016).

In all species, CSL is predominantly nuclear even though it lacks a recognizable nuclear localization signal, making it likely that other factors chaperone CSL into the nucleus. NICD and several co-repressors, such as Hairless and SMRT, can fulfil this role and promote nuclear accumulation of CSL (Fortini and Artavanis-Tsakonas, 1994; Gho et al., 1996; Maier, 2020; Maier et al., 2013; Tabaja et al., 2017; Wacker et al., 2011; Wolf et al., 2019, 2021; Zhou and Hayward, 2001). However, it has been widely assumed that NICD first meets CSL in the nucleus, although there is no direct evidence to support this, and it remains possible that two proteins could interact prior to nuclear entry. The main argument against is that CSL does not interact with the full-length transmembrane Notch receptor, despite the fact that NICD would be exposed in the cytoplasm (e.g. Gho et al., 1996). To investigate the relationship between NICD release and CSL transport into the nucleus, we developed OptIC-Notch, an optogenetic tool in which NICD is tethered to the membrane and released upon light activation.

## Results and Discussion

### Membrane tethered NICD can sequester CSL away from the nucleus

OptIC-Notch uses the BLITz approach (Lee et al., 2017) in which light exposure simultaneously reconstitutes a TEV protease and reveals a TEV target site in the membrane tethered protein, resulting in its release. In this case we used two constructs. In the first, a CIBN domain, N-terminal TEV protease fragment and a TEV cleavage-site motif were inserted between NICD fused with mCherry and a heterologous transmembrane domain (CIBN-TEVn-mCherry-NICD, Fig. 1A). In the second, a cytoplasmic Cryptochrome 2 photolyase homology region was fused to a C-terminal TEV protease fragment (CRY-TEVc). When co-expressed, the active TEV protease will be reconstituted by light induced association of CIBN and CRY domains and can cleave at the adjacent TEV cleavage motif to release mCherry::NICD. As in the original BLITz, the TEV motif is protected by a light sensitive AsLOV2 domain (LOV) to prevent leaky cleavage (Fig. 1A; (Lee et al., 2017)).

**Figure 1:**
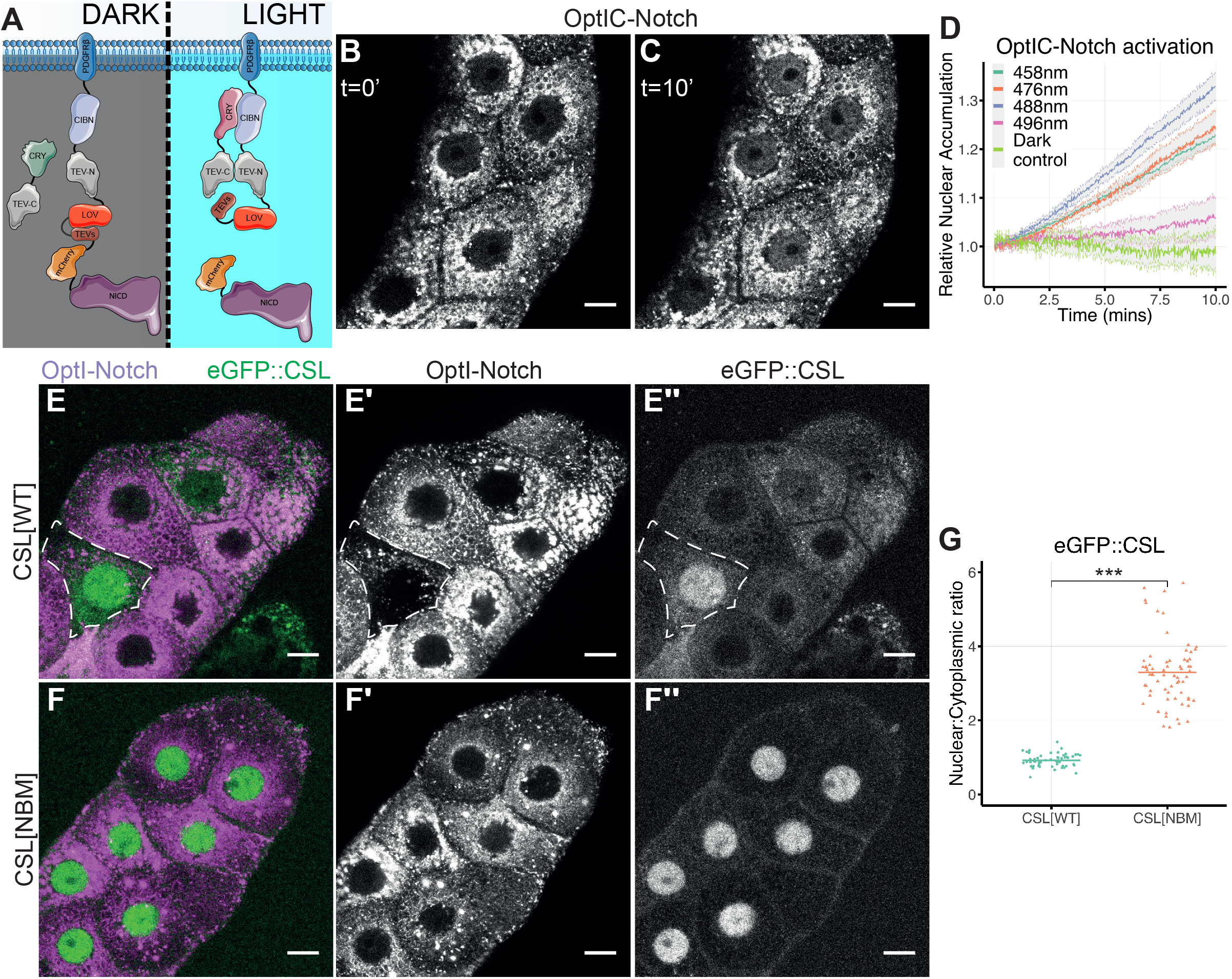
CSL is sequestered from the nucleus by OptIC-Notch. (A) Schematic depicting domain structure of OptIC-Notch and blue light induced release of NICD (purple). Blue light induces (i) heterodimerisation of CRY2 (green) and CIBN (lilac) domains, reconstituting TEV protease (TEV-C and TEV-N; grey) (ii) unfolding of LOV (Red) reveals TEV cleavage site (TEVs, Brown), cutting releases NICD. Note that in some experiments mCherry was replaced by GFP. (B-C) Localization of OptIC-Notch in salivary glands before (B; t=0’) and after 10 minutes in 458nm (blue) light (C, t=10’); mCherry::NICD is detected in the nucleus following light exposure. Scale bar = 20 μm. (D) Relative nuclear accumulation, normalised to t=0 min, of mCherry::NICD after exposure of salivary glands expressing OptIC-Notch to indicated wavelengths of light over a 10 minute period. Dark controls were not exposed to light (mean ± SEM). (E) Localisation of eGFP::CSL (green, E, single channel E’’) in salivary glands expressing OptI-Notch (magenta, E, single channel E’;). eGFP::CSL is sequestered outside the nucleus in most cells; dashed outline, cell with low levels of OptI-Notch and higher nuclear eGFP::CSL. Scale bar = 20 μm. (F) eGFP::CSL[NBM] (green, F, single channel F’’) is not sequestered by OptI-Notch (magenta, F, single channel F’), conditions as in E. Scale bar = 20 μm. (G) Nuclear to cytoplasmic ratio of eGFP::CSL[WT] and eGFP::CSL[NBM] in the presence of OptI-Notch (without CRY-TEVc). Bar represents mean of ≥10 salivary glands per condition. Welch’s two tailed t-test p-value < 0.001.

Expressing the trans-membrane construct in isolation will not support cleavage. We refer to this uncleavable condition as OptI-Notch and to the cleavable combination as OptI<u>C</u>-Notch. Transgenic flies were generated to produce OptI-Notch and OptIC-Notch under control of tissue-specific UAS-Gal4 combinations.

When expressed in *Drosophila* third instar larval salivary glands in the dark, OptIC-Notch was present in membranous structures throughout the cytoplasm and absent from the nucleus. Exposure to blue light resulted in rapid nuclear accumulation of cleaved NICD (Fig. 1B-D, Sup MOV 1). Consistent with the dependence on blue light induced cleavage, the accumulation was weaker with longer wavelengths, that would be less effective at reconstituting TEV (496 nm) and was non-existent in dark control conditions, where the salivary glands were not exposed to blue light (Fig. 1D). These results confirm that NICD can be released from OptIC-Notch in a light dependent manner.

To investigate the interactions between NICD and CSL, we expressed OptI-Notch/OptIC-Notch in the context of fluorescently tagged CSL (eGFP::CSL or Halo::CSL) present at endogenous levels (Gomez-Lamarca et al., 2018). Strikingly, in tissues expressing OptI-Notch, eGFP::CSL was strongly enriched in the cytoplasm with little or none present in the nucleus (Fig. 1E and F). In the cytoplasm, CSL colocalized with the OptI-Notch suggesting that it was becoming sequestered by interacting with the membrane tethered NICD. To confirm that CSL sequestration was mediated by its interaction with NICD, we performed the same experiment with a mutated version of CSL defective in Notch-binding (CSL[NBM] (Gomez-Lamarca et al., 2018; Yuan et al., 2016)). CSL[NBM] remained localised to the nucleus in the presence of OptI-Notch (1E-G). These data argue that CSL is sequestered by an interaction with the NICD moiety in OptI-Notch. Consistent with this hypothesis a different transmembrane tethered NICD, in which mCherry-NICD was fused to a CD8 transmembrane domain, also sequestered a significant fraction of CSL in the cytoplasm (Sup Fig. S1).

### Released NICD chaperones CSL into the nucleus

Next, we investigated whether sequestered CSL could be effectively transported into the nucleus when NICD was released from the membrane by exposing larvae expressing OptIC-Notch to blue light for 24 hours. Under these conditions there was a clear change in CSL localisation, with a significant fraction relocating to the nucleus (Fig. 2A-C). These results suggest that, having interacted with the membrane tethered NICD in the cytoplasm, CSL can be carried into the nucleus by NICD when it is released.

**Figure 2:**
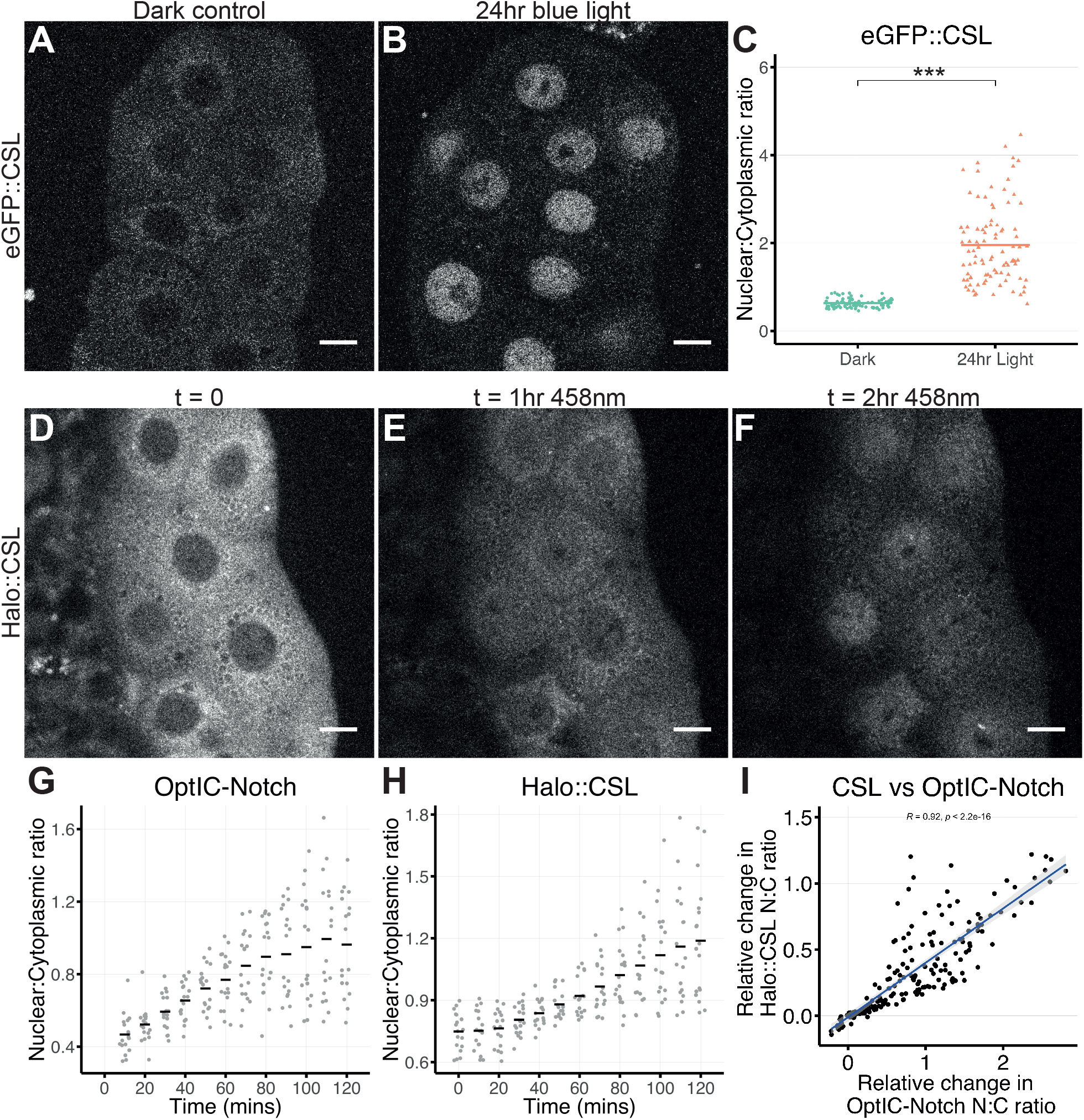
Relocalization of CSL to the nucleus when NICD is released. (A,B) Localisation of eGFP::CSL in salivary glands expressing OptIC-Notch without activation (A) and after 24hrs of blue-light induced activation (B). Cytoplasmic sequestration of eGFP::CSL is reversed. Scale bar = 20 μm. (C) Nuclear to cytoplasmic ratio of eGFP::CSL is increased after 24 hours of blue-light exposure in a custom incubator, bar represents mean of >10 salivary glands per condition. Welch’s two tailed t-test p-value < 0.001. (D-F) Localization of Halo::CSL in the presence of OptIC-Notch (GFP variant) detected before (t=0) and after exposure to continuous blue light (458 nm) for indicated times. Salivary glands were exposed to Halo ligand JF646 prior to imaging. By 2hrs a significant fraction of Halo::CSL has translocated to the nucleus. Scale bar = 20 μm. (G-H) Light exposure leads to increased nuclear:cytoplasmic ratio of OptIC-Notch (G, indicative of NICD release) and of Halo::CSL (H). (I) Rate of nuclear accumulation of GFP::NICD and of Halo::CSL. The two are correlated (Spearmans test, Rho=0.92, p < 0.001).

To rule out the possibility that the nuclear CSL is newly translated protein, which enters the nucleus independently and becomes stabilised by the released NICD, we performed a pulsed labelling with the Halo ligand JF646. Salivary glands expressing Halo::CSL and OptIC-Notch were incubated with JF646 and subsequently washed to remove unbound ligand. Any CSL synthesized subsequently would thus be unlabelled. Cleavage of OptIC-Notch was then induced by blue light exposure whilst at the same time imaging JF646 bound Halo::CSL with far-red illumination. After two hours a clear nuclear accumulation of CSL was detected (Fig. 2D-F). The nuclear levels of NICD (Fig. 2G) and Halo::CSL (Fig. 2H) increased over time and the latter were proportional to the amount of nuclear NICD detected (Fig. 2I). Thus, the nuclear CSL corresponds to protein that was previously sequestered in the cytoplasm and that was likely transported to the nucleus by the released NICD.

We next asked whether NICD that is released by γ-secretase cleavage (Kopan and Ilagan, 2009) is capable of transporting CSL into the nucleus. Because the co-repressor Hairless has an important role in transporting and stabilising CSL in the nucleus (Fechner et al., 2022; Praxenthaler et al., 2017), we first used RNAi to knock down expression of Hairless to avoid confounding effects. These conditions resulted in a decrease in overall levels of nuclear CSL (Sup Fig. S2 A,B), as described previously (Gomez-Lamarca et al., 2018; Praxenthaler et al., 2017). When combined with a transgene producing constitutively cleaved Notch (Nact), the levels of nuclear CSL were largely restored, suggesting that released NICD is able to transport CSL into the nucleus (Sup Fig. S2C,D,E; and see Gomez-Lamarca et al., 2018; Praxenthaler et al., 2017). Altogether, the results demonstrate that CSL and NICD can interact in the cytoplasm and are co-transported into the nucleus. However, this raises the question why CSL does not normally interact with the full-length endogenous Notch receptor before its cleavage (Gho et al., 1996). This lack of colocalization is evident in the epithelial follicle cells of the Drosophila egg chamber, where the Notch receptor is enriched at the apical and lateral surfaces with no CSL associated (Sup Fig. S3). However, in a few cells we detected cytoplasmic and perinuclear puncta where the two proteins were colocalized suggesting they can associate outside the nucleus under certain circumstances (Sup Fig. S3C).

### Masking ΦWΦP motif prevents CSL interaction

One hypothesis to explain why CSL is sequestered by OptIC-Notch but not by the endogenous receptor at the membrane, is that the high affinity CSL-binding ΦWΦP motif (Fig 3A; (Choi et al., 2012; Hall and Kovall, 2019; Johnson et al., 2010; Kovall and Hendrickson, 2004; Lubman et al., 2007; Nam et al., 2003; Wilson and Kovall, 2006) is exposed in OptIC-Notch but is normally hidden. It has been proposed that, in its normal context, this hydrophobic motif, which is located close to the transmembrane domain, becomes embedded in the lipid bilayer of the plasma membrane (Deatherage et al., 2015, 2017). This would occlude it from CSL until NICD is released from the membrane by γ-secretase cleavage. The insertion of several domains between the membrane and the ΦWΦP motif in OptIC-Notch could change the protein conformation, preventing the association of the motif with the membrane and making it available for binding with CSL.

**Figure 3:**
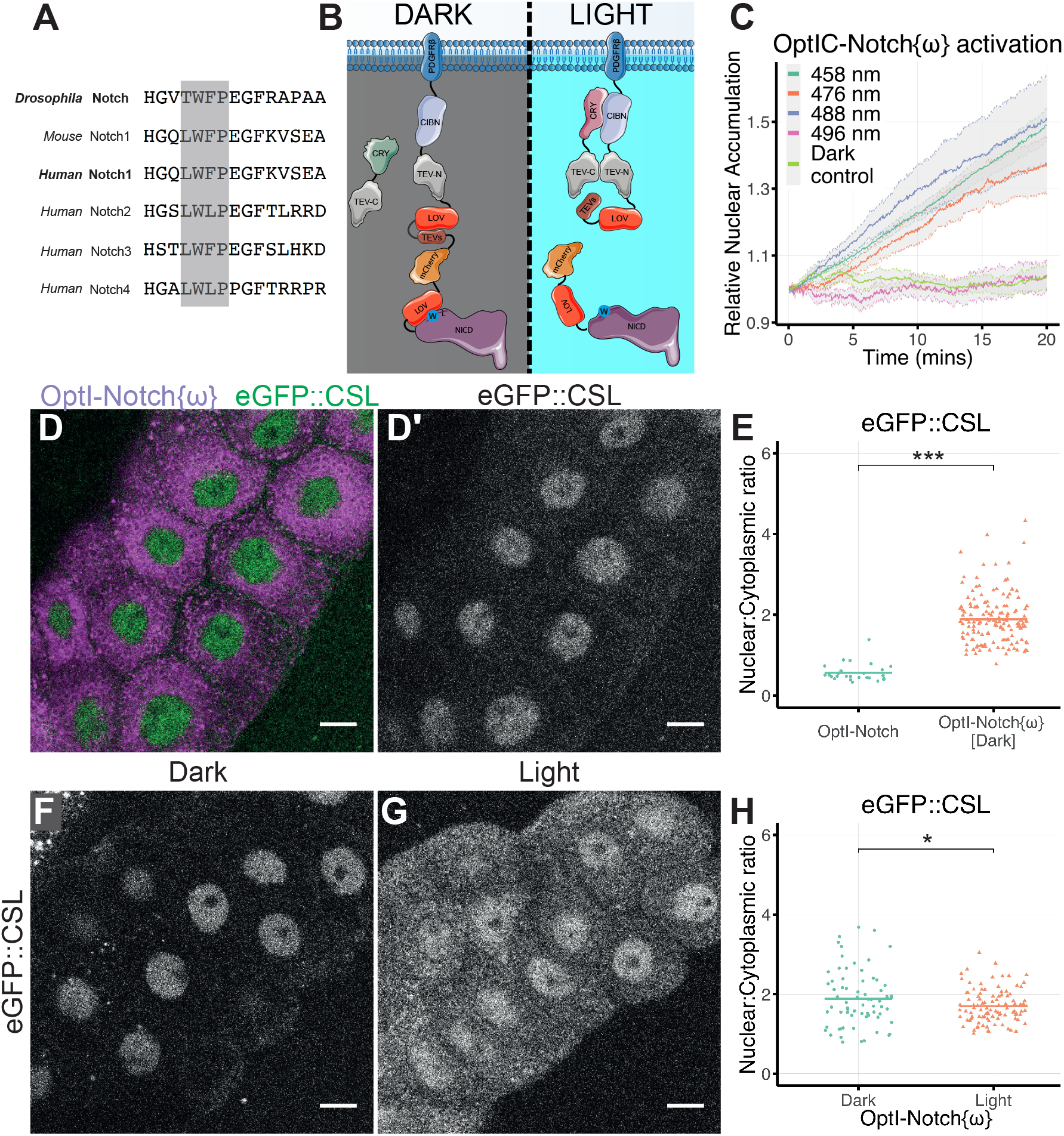
Masking ΦWΦP motif prevents CSL sequestration. (A) Sequence alignment of residues from the indicated NICD with the conserved ΦWΦP motif highlighted (grey shading). (B) Schematic of modified optogenetic construct, OptIC-Notch{ω}, containing an additional LOV domain (red) to mask the ΦWΦP motif (blue, W) in NICD. Other domains are as in Fig. 1A. (C) Relative nuclear accumulation, normalised to t=0 min of mCherry::NICD after exposure of salivary glands expressing OptIC-Notch{ω} to indicated wavelengths of light over 20 minutes. Dark controls were not exposed to light (mean ± SEM). (D) eGFP::CSL (D, green; D’) is nuclear in the presence of OptI-Notch{ω} without CRY-TEVc (D, magenta). (E) Nuclear to cytoplasmic ratio of eGFP::CSL in tissues expressing OptI-Notch{ω} in comparison to OptI-Notch (both without CRY-TEVc). Significantly higher ratios are detected with OptI-Notch{ω}. Bar represents mean of ≥5 salivary glands per condition. Welch’s t-test p-value < 0.001. (F-H) Localization of eGFP::CSL in tissues expressing OptI-Notch{ω} (without CRY-TEVc) in dark (F) and under blue light exposure in a custom incubator (G), the nuclear to cytoplasmic ratio is reduced (H). In (H) Bar represents mean of > 10 salivary glands per condition. Welch’s two tailed t-test p-value = 0.048. Scale bars in D,F,G = 20 μm.

If this hypothesis is correct, CSL should no longer be recruited by OptIC-Notch when the ΦWΦP motif is masked. To achieve this, we added a second LOV domain to OptIC-Notch, six residues upstream of the ΦWΦP motif in NICD, here referred to as OptIC-Notch{ω} (Fig. 3B). In the dark, the additional LOV domain in OptIC-Notch{ω} will conceal the ΦWΦP motif. If this motif is responsible for the interaction, CSL should no longer interact with the NICD tethered to the membrane. Upon blue light activation OptIC-Notch{ω} is cleaved in the same way as OptIC-Notch and, at the same time, the LOV domain concealing the ΦWΦP motif will unfold to make this motif available for CSL binding (Fig. 3B-C).

To assess the effects on CSL, we first expressed the membrane bound construct without the cytoplasmic CRY-TEVc (OptI-Notch{ω}). When kept in the dark, so that the ΦWΦP motif was protected by the LOV domain, CSL was no longer sequestered as predicted if the accessibility of the motif is a critical factor (Fig. 3D). Indeed, under these conditions the majority of CSL was detected in the nucleus (Fig. 3E). Furthermore, expression of OptI-Notch{ω} in the developing wing produced no discernible phenotype, unlike expression of the parent OptI-Notch which resulted in malformed adult wings with blisters and aberrant vein patterning likely due to reduced availability of CSL for endogenous Notch signalling (Sup Fig. S4).

Exposure of OptI-Notch{ω} to light should unmask the ΦWΦP motif, by unfolding the LOV domain, and, in the absence of CRY-TEVc, NICD will be tethered because no cleavage would occur. In these conditions a significant fraction of CSL became sequestered with the OptI-Notch{ω} outside the nucleus, consistent with the exposure of the ΦWΦP motif being a critical factor in its relocalization (Fig. 3F-G). The sequestration was not as robust as that with the parent construct, and some CSL remained in the nucleus. Likely explanations are that a fraction of the CSL was retained there by interactions with nuclear proteins such as Hairless and/or that unmasking was not 100% effective. Nevertheless, the fact that masking of the ΦWΦP motif largely eliminates CSL sequestration, and its unmasking partially reverses that, argues that this motif is responsible for the interaction. Furthermore, it is evident that the motif must be hidden in the native conformation of the endogenous receptor to prevent CSL sequestration. As its exposure upon cleavage is a key feature for promoting formation of CSL-NICD complex, it’s possible that this interaction could occur prior to nuclear entry.

### Light induced NICD release mimics Notch signalling

A final question is whether light induced release of NICD from the membrane is competent to activate Notch signalling. As with OptIC-Notch, blue light activation of OptIC-Notch{ω} resulted in a rapid nuclear accumulation of NICD, indicative of efficient cleavage (Fig. 4A-C and 3C, Sup MOV 2). This did not occur with longer wavelengths of light that cannot induce the reconstitution of the TEV. Indeed, two hours of light were sufficient to bring about robust nuclear enrichment of NICD (Fig. 4B-C) and to promote the recruitment of CSL to the target *E(spl)-C* locus (Fig. 4D-F). This was visualized by the presence of an enriched band of CSL that co-localized with the fluorescent ParB-Int labelled genomic *E(spl)-C* locus, as described previously (Fig. 4E-F; (Gomez-Lamarca et al., 2018). Furthermore, robust expression of the Notch target gene *E(spl)mβ* was detected in blue light conditions, demonstrating that the released NICD was functional and able to induce target gene transcription (Fig 4G-I).

**Figure 4:**
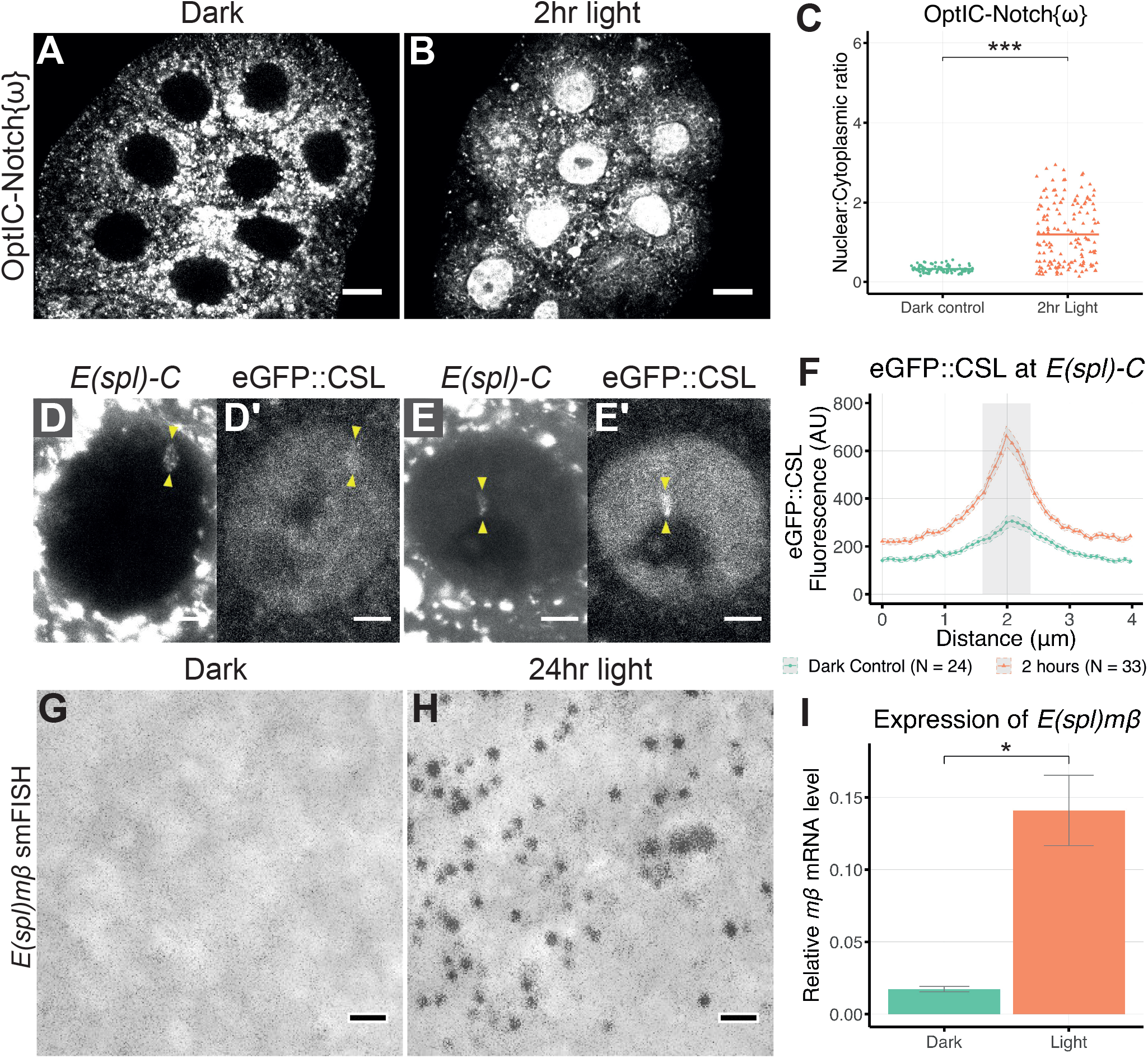
Effects of light induced cleavage of OptIC-Notch{ω} on CSL localization and *E(spl)mβ* expression. (A-B) Localization of OptIC-Notch{ω} before (A) and after 2hrs of continuous blue light in a custom incubator (B). Strong nuclear localisation of mCherry::NICD in B is indicative of efficient light induced cleavage. Scale bars = 20 μm. (C) Nuclear to cytoplasmic ratio of mCherry::NICD in salivary glands kept in the dark and 2hrs light, as in 4B. Bar represents mean of >10 salivary glands. Welch’s T-test p value < 0.001. (D-F) Nuclear distribution of eGFP::CSL in the presence of OptIC-Notch{ω}. No/little enrichment at *E(spl)-C* is detected in dark conditions (D’,F), strong enrichment at *E(spl)-C* is detected within 2hrs of light, as in 4B (E’,F). *E(spl)-C* is visualized using parB-Int system in D, E. The corresponding region is depicted by grey shading in F in relation to the quantified eGFP fluorescence intensity. Scale bars = 5 μm, error bars = ± SEM. (G-H) Detection of cytoplasmic *E(spl)mβ* transcripts by smFISH (fluorescent puncta, black) in salivary glands kept in the dark (G) or after 24hr continuous blue light in a custom incubator (H), fluorescent images are overlaid on transmitted light, scale bar = 1 μm. (I) Quantification of *E(spl)mβ* RNA by RT-qPCR in the same two conditions shows significant increase, N=3, error bars are ± SEM. Welch’s two tailed t-test p-value = 0.036.

## Conclusion

In conclusion, our optogenetic tools to control release of NICD have highlighted the importance of masking the ΦWΦP motif to prevent sequestration of CSL by the transmembrane receptor. When this motif is exposed, CSL interacts with NICD in the cytoplasm. In our constructs, the masking is achieved with a light-regulated LOV domain. In the native receptor it likely relies on shielding by association with membrane phospholipids (Deatherage et al., 2015, 2017). Our findings are fully consistent with the structural studies which have shown the importance of the RAM domain, containing the ΦWΦP motif, for binding to CSL and suggest the mechanism will be conserved across species (Choi et al., 2012; Contreras et al., 2015; Friedmann et al., 2008; Hall and Kovall, 2019; Nam et al., 2003; Wilson and Kovall, 2006). Previous studies have emphasized that this motif has a role in the nucleus. Here we suggest that, as it will be exposed immediately upon cleavage, CSL could partner with NICD in the cytoplasm. In this way, CSL would be transported into the nucleus with NICD after forming a complex. Tight regulation of domain-motif interactions is of fundamental importance for many central cellular processes, and frequently relies on post-translational modifications (Akiva et al., 2012). The membrane masking of the CSL binding-motif in Notch is an interesting example of such regulation.

Although our experiments are in *Drosophila*, the ΦWΦP motif and its interaction with CSL is highly conserved (Choi et al., 2012; Contreras et al., 2015; Hall and Kovall, 2019; Johnson et al., 2010; Nam et al., 2003; Wilson and Kovall, 2006). Likewise, shuttling of CSL between nuclear and cytoplasmic compartments has been observed in several contexts, where its regulation appears important for fine-tuning Notch dependent transcription (Fechner et al., 2022; Gho et al., 1996; Maier, 2020; Maier et al., 2013; Tabaja et al., 2017; Wacker et al., 2011; Wolf et al., 2019, 2021; Zhou and Hayward, 2001). Whether these impacts on signalling are because of consequences on nuclear CSL pools, which has largely been the inference, or whether effects on cytoplasmic pools are also important as implied here, remains to be established. Our results, that CSL can be imported into the nucleus by NICD after first meeting in the cytoplasm, make it plausible that the balance of cytoplasmic CSL will be an important facet in Notch regulation.

### Caveats to the approach

One caveat with our approach is that it relies on over-expressed synthetic Notch proteins although we also detect occasional cytoplasmic puncta where CSL and Notch colocalize in normal conditions. Genome engineered endogenous proteins with altered motif exposure would avoid the technical problems from over-expressed proteins but, as they would dominantly sequester CSL, it is likely they would cause lethality and be unrecoverable. It therefore remains to be proven whether CSL and NICD normally interact outside the nucleus under physiological conditions.

## Acknowledgements

We thank Kat Millen and the Genetics Department Fly Facility for performing DNA injections of fly embryos to generate transgenic stocks. We are grateful to Cambridge Advanced Imaging Centre, and all members of the Bray Lab, for helpful discussions. We also acknowledge Servier Medical Art, for images used in the generation of figures 1A and 3B.

## Author contributions

JMT, MGL and SJB designed the experiments. JMT and CSCM performed the experiments. MGL generated critical reagents. JMT and SJB analysed the data. JMT and SJB wrote the manuscript, MGL reviewed and commented on the manuscript.

## Townson et al; Supplementary Material

**Supplementary Movie 1: Release of NICD from OptIC-Notch**.

OptIC-Notch is activated with 458 nm light for 10 minutes. Release of NICD and its accumulation in the nucleus over time is shown by simultaneous imaging of the mCherry tag. Scale bar = 20 μm.

**Supplementary Movie 2: Release of NICD from OptIC-Notch{ω}**

OptIC-Notch{ω} is activated with 458 nm light for 20 minutes. Release of NICD and its accumulation in the nucleus over time is shown by simultaneous imaging of the mCherry tag. Scale bar = 20 μm.

**Supplementary Figure S1:**
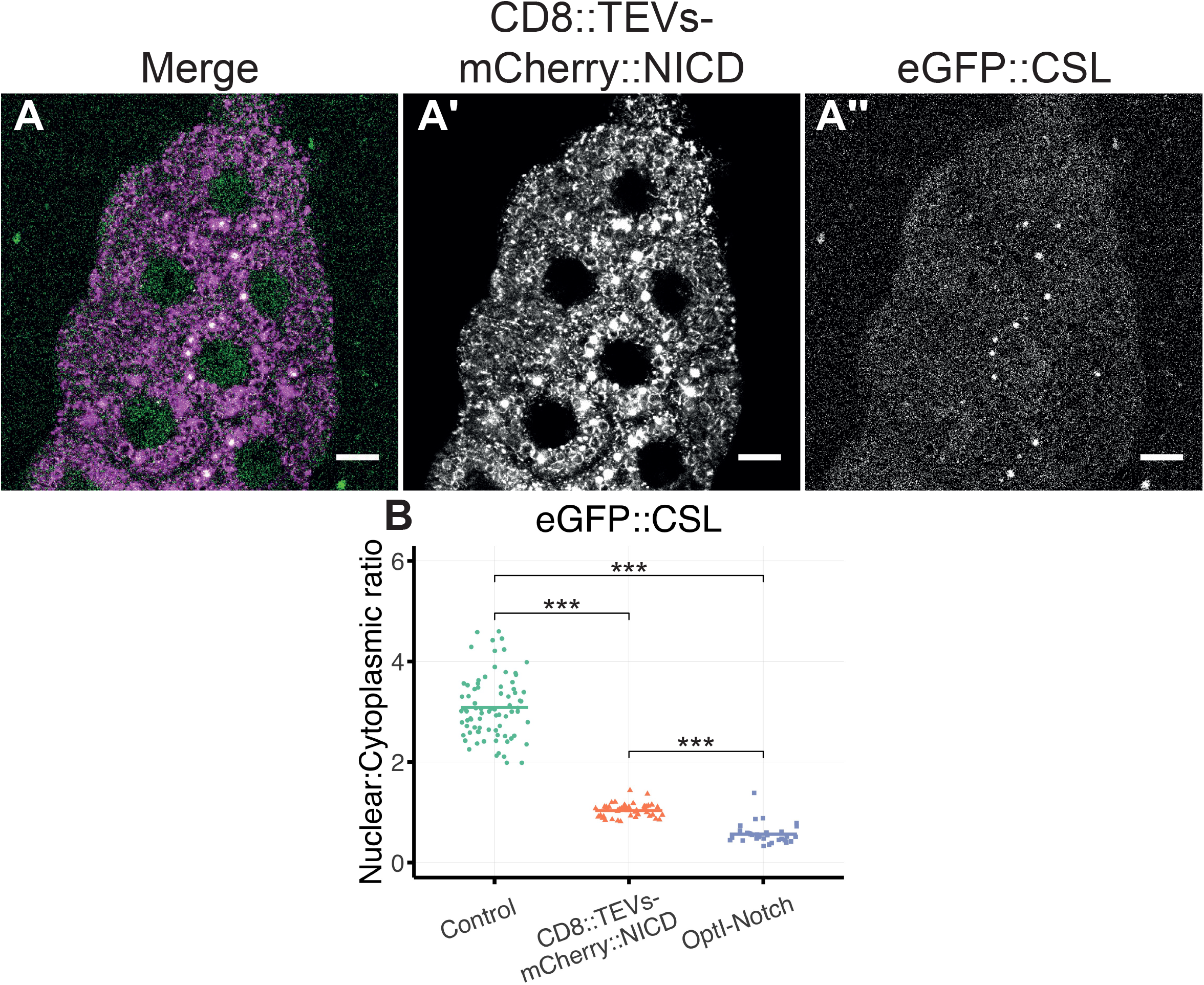
CSL is sequestered by additional constructs containing NICD. (A) Localisation of eGFP::CSL in salivary glands expressing a construct containing a CD8 transmembrane domain fused to NICD, with a TEV site and mCherry tag in between (magenta and A’). CSL is partially sequestered in the cytoplasm (green and A’’). Scale bar = 20 μm. (B) Nuclear to cytoplasmic ratio of eGFP::CSL in the presence of the CD8 tethered construct and OptI-Notch (without CRY-TEVc), as a control condition, data from flies expressing LacZ and *white* RNAi were used, as shown in Fig S2. Bar represents the mean of ≥7 salivary glands per condition. Welch’s two tailed t-test p-value < 0.001.

**Supplementary Figure S2:**
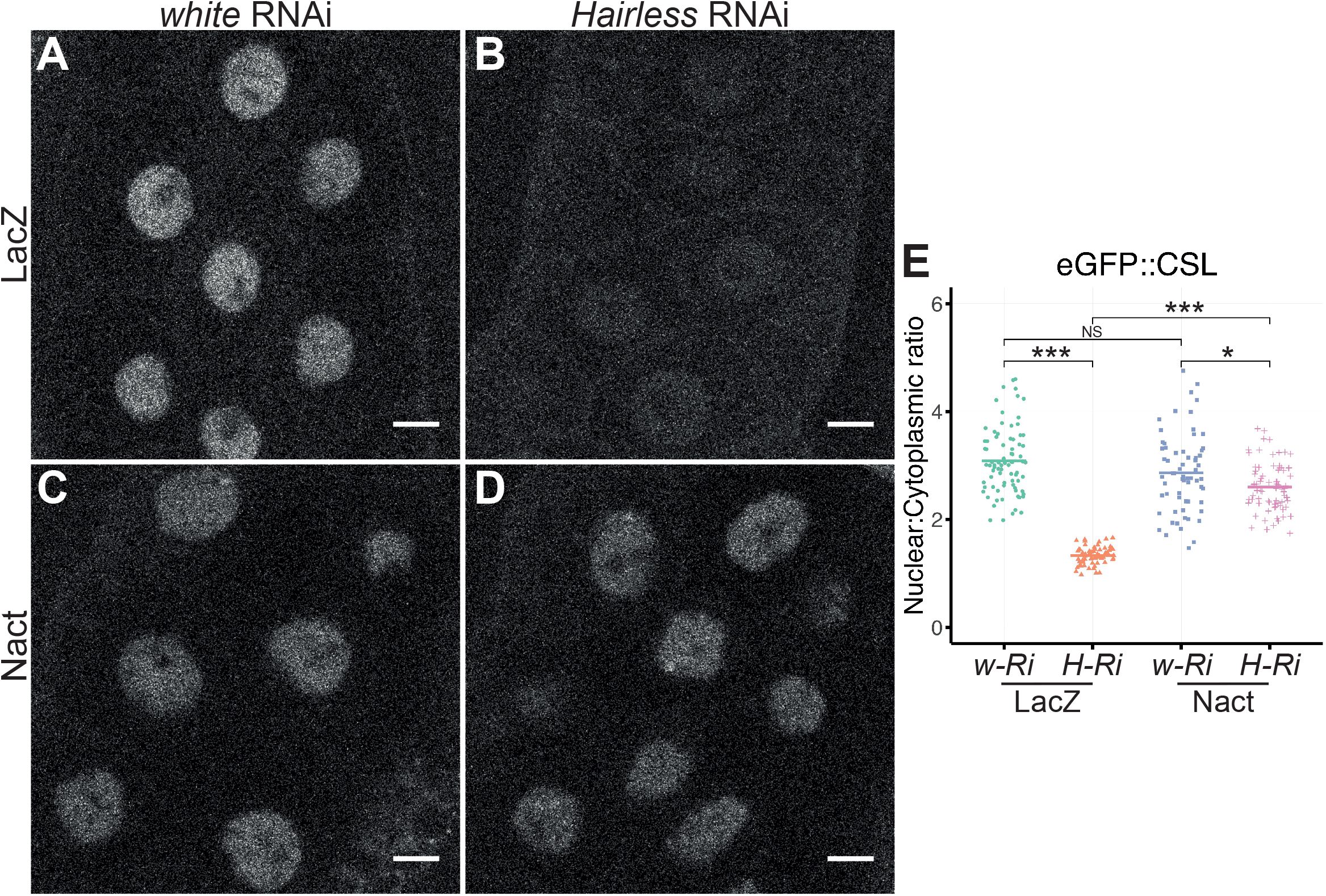
Effects of Notch on nuclear levels of CSL in the absence of Hairless. **(A-D)** Localisation of eGFP::CSL in salivary glands expressing LacZ or Nact (the constitutively active Notch, NΔECD) with *white* RNAi (A, C) or *Hairless* RNAi (B, D). A large decrease in CSL levels is seen in the absence of Hairless (B) This decrease in CSL levels is not seen with Nact in the absence of Hairless (D). Scale bar = 20 μm. **(E)** Nuclear to cytoplasmic ratio of eGFP::CSL in tissues expressing LacZ or Nact in a *white* or *Hairless* RNAi background. Bars represent the mean of >10 salivary glands per condition. Welch’s two tailed t-test p-values for LacZ *w-Ri* vs *H-Ri* < 0.001, Nact *w-Ri* vs *H-Ri* = 0.012, *H-Ri* LacZ vs Nact < 0.001.

**Supplementary Figure S3:**
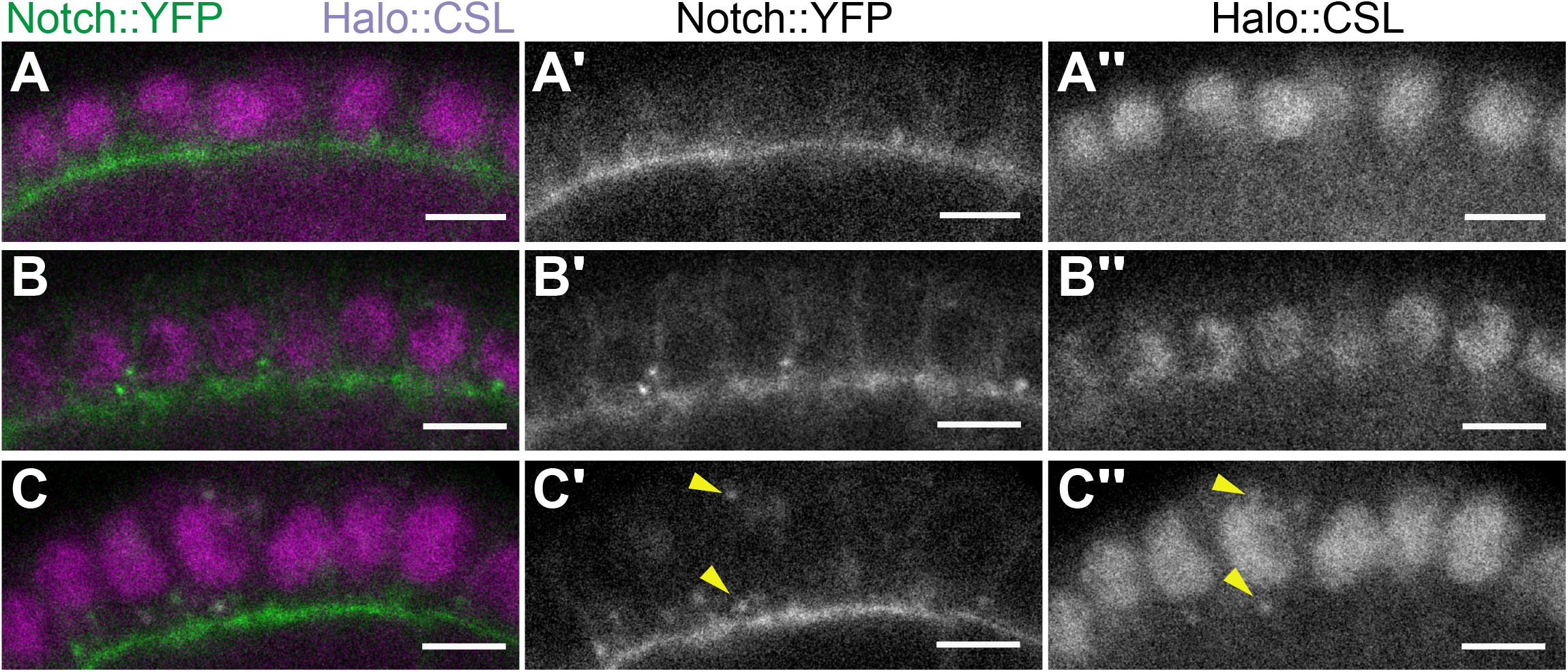
Notch and CSL do not colocalize at the cell membrane. **(A-C)** High-resolution confocal images of stage six egg chambers from *Drosophila* ovaries showing Notch::YFP (A’-C’) and CSL::Halo (A’’-C’’) in the follicle cells. Notch decorates apical and lateral membranes of follicle cells, whereas CSL is restricted to the follicle cell nuclei. In a few cases, co-localization is observed in cytoplasmic puncta (C, yellow arrows in C’ and C’’). Scale bars = 5 μm.

**Supplementary Figure S4:**
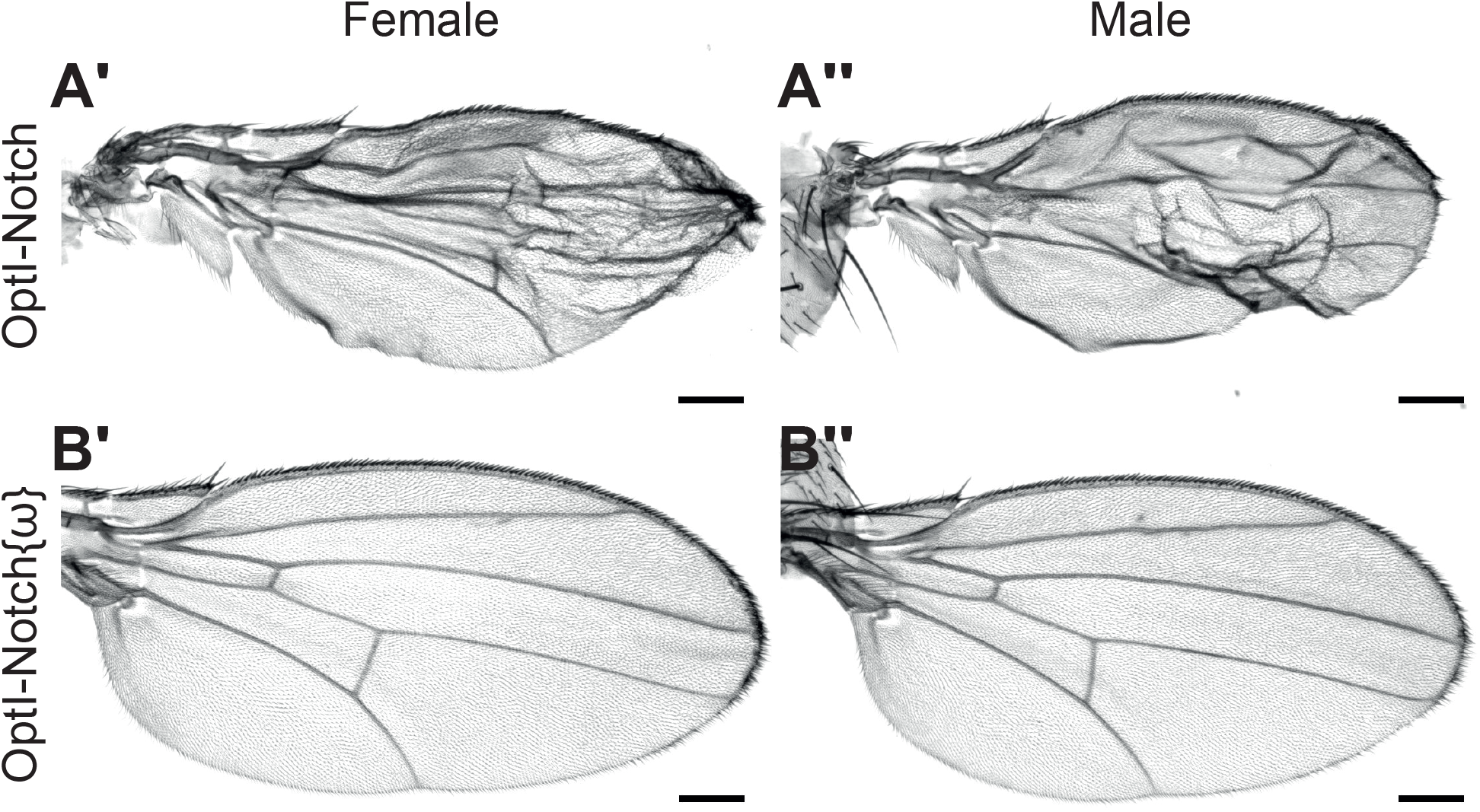
OptI-Notch{ω} does not generate phenotypes in wings seen with OptI-Notch. OptI-Notch (A) or OptI-Notch{ω} (B) is expressed in the wing in both males and females. Blistering and aberrant vein patterning occurs with OptI-Notch and not OptI-Notch{ω}. Scale bar = 100 μm.

## Materials and Methods

### RESOURCE AVAILABILITY

#### Lead contact

Further information and requests for resources and reagents should be directed to, and will be fulfilled by, the Lead Contact, Sarah J Bray.

#### Materials availability

All stocks, plasmids, and reagents generated in this study are available from the Lead Contact but we may require a completed Materials Transfer Agreement.

#### Data and code availability

Microscopy data reported in this paper will be shared by the lead contact upon request. All original code has been deposited at Zenodo and is publicly available as of the date of publication. DOIs are listed in the key resources table.

Any additional information required to reanalyze the data reported in this paper is available from the lead contact upon request.

### Key Resources Table

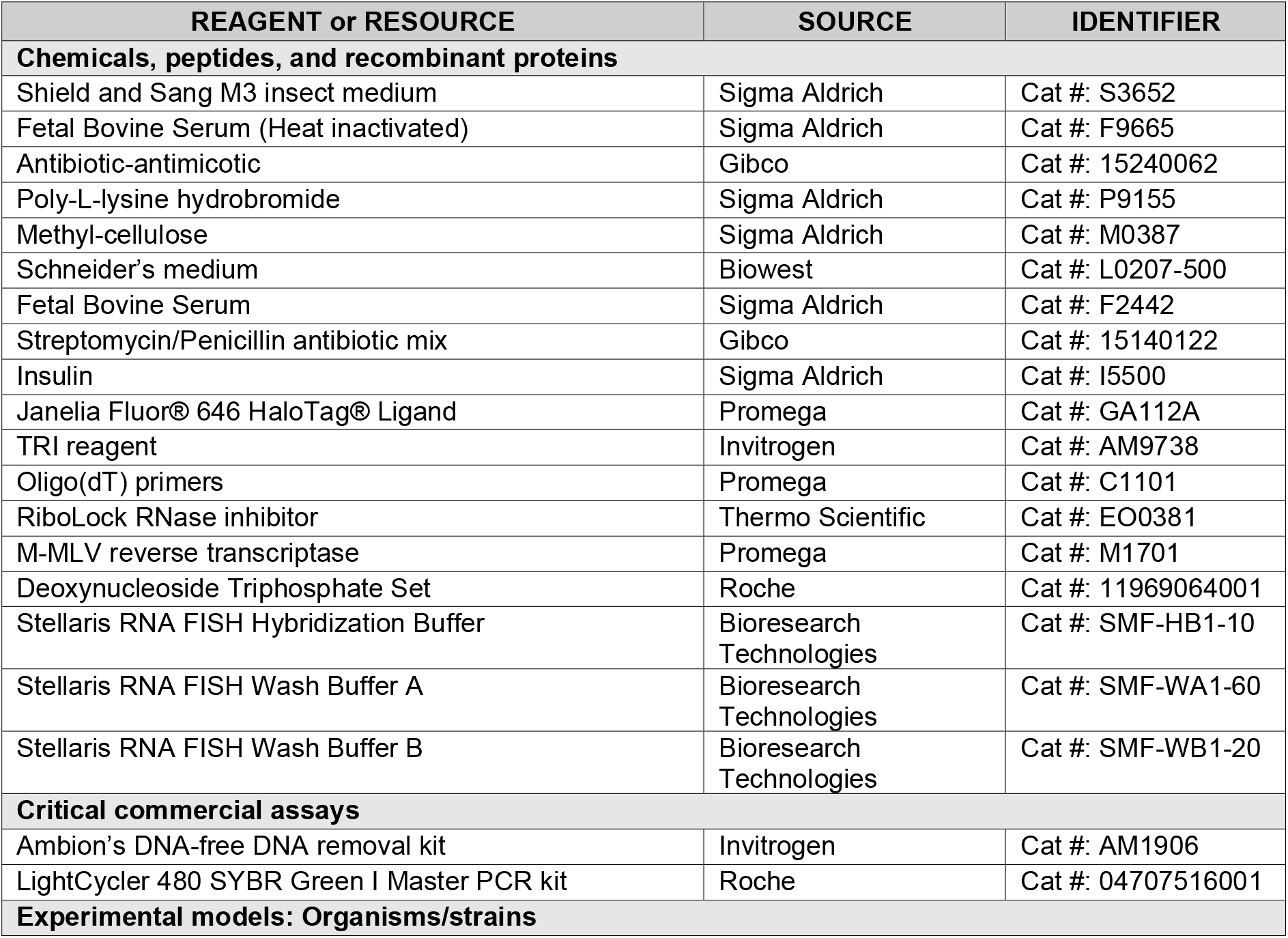
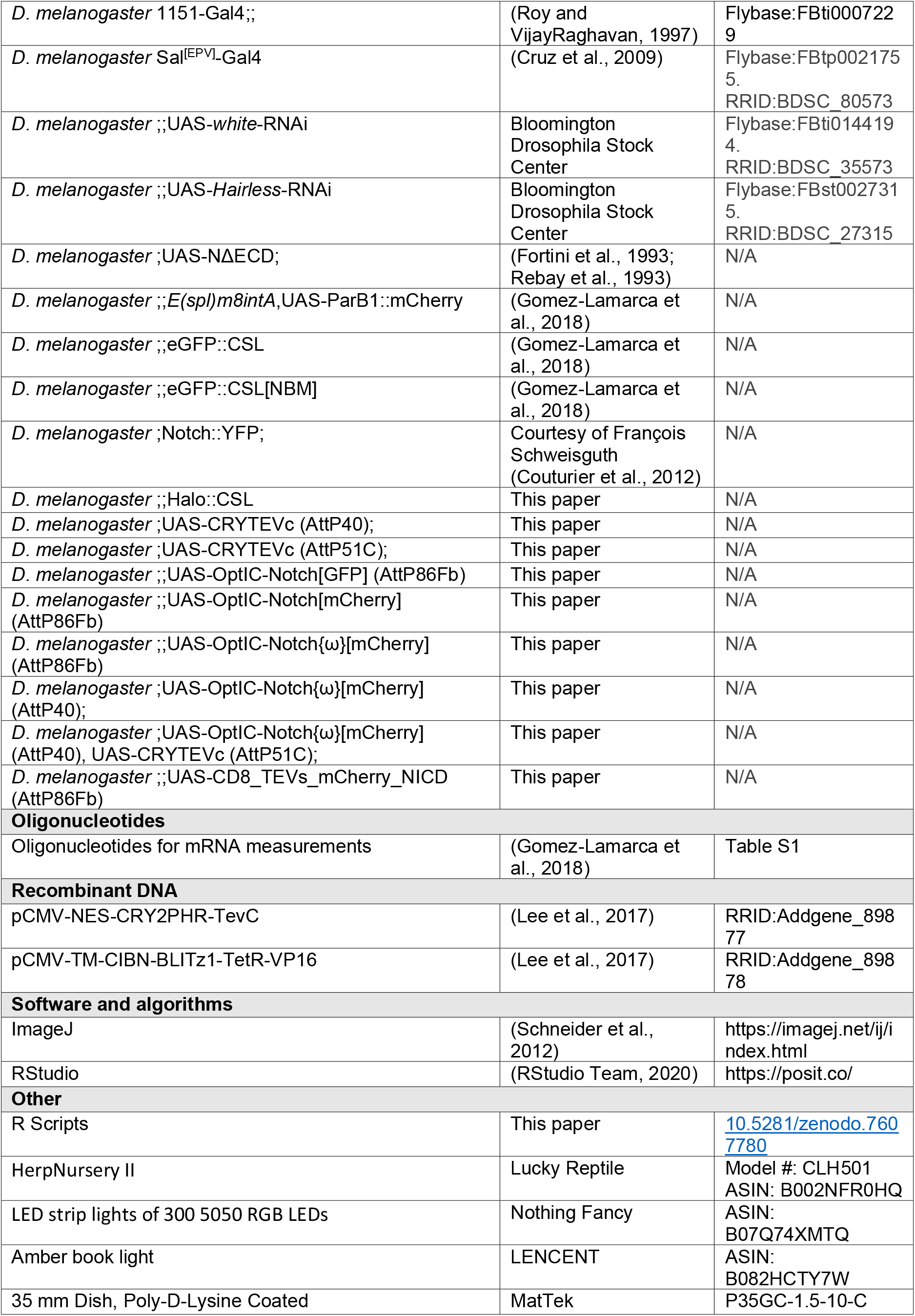

## EXPERIMENTAL MODEL AND CONDITIONS

### *Drosophila melanogaster* strains and genetics

All *Drosophila melanogaster* stocks were grown on standard medium at 25 °C. Details of stocks used are available in the key resources table. Expression from the indicated constructs was driven using *1151-Gal4*, which gives robust expression in the salivary glands as well as in the muscle precursors (Roy and VijayRaghavan, 1997) and *Sal*^*[EPV]*^*-Gal4* containing a fragment from Salm that directs expression in the central pouch region of 3^rd^ instar wing discs (Cruz et al., 2009). Endogenous levels of Notch and CSL were imaged in egg chambers using a YFP tagged Notch construct (Couturier et al., 2012).

For dark conditions, vials were wrapped in aluminium foil and kept in a thick cardboard box. For global blue light illumination over extended periods of time, a blue light incubator was built as in (Grusch et al., 2014). Briefly, an incubator with temperature control (LuckyReptile, HerpNursery II) was fitted with LED strip lights of 300 5050 RGB LEDs and set to constant blue illumination. Larvae were placed in the incubator on apple juice plates supplemented with yeast, for maximal light exposure, according to the length of time specified. For the inducible sequestration experiments larvae were grown in the light from embryos laid on apple juice plates and corresponding dark controls were grown the same way, wrapped in foil inside a dark cardboard box.

### Generation of optogenetic fly lines and Halo::CSL

BLITz constructs (Lee et al., 2017) were obtained from Addgene (89877 and 89878) and the tTA domain replaced with an NICD fragment (residues T1766 to I2703 of the full-length *Drosophila* Notch receptor, Uniprot P07207), fused with mCherry or GFP at the N terminus, to generate OptIC-Notch. The second LOV domain in OptIC-Notch{ω} was added by Gibson cloning between position R1768 and K1769 (numbering of the full-length Notch receptor) to the NICD fragment of OptIC-Notch. The CD8-TEVs-mCherry-NICD construct was cloned by the replacement of the BLITz element in OptIC-Notch with CD8-TEVs using restriction digest cloning. All constructs were then cloned into a UASt vector containing AttB integration sites (Bischof et al., 2007).

Transgenic optogenetic fly lines were generated by injection of plasmid DNA into embryos and integration by germ-line specific ΦC31 integrases (Bischof et al., 2007) into AttP sites on the second and third chromosome (AttP40, AttP51C, AttP86Fb). Male transgenics were selected using miniwhite.

A Halo::CSL construct expressed at endogenous levels was generated as in (Gomez-Lamarca et al., 2018), using a Halo tag in place of the N-terminal GFP tag. Labelling of the Halo tag in salivary glands and ovaries was performed by incubating in the respective dissection media containing 1 μM (salivary glands) and 0.1 μM (ovaries) of JF646 ligand (Promega, GA112A) for 15 minutes, followed by three 15-minute washes in dissection media.

### Sample dissection and mounting

Salivary glands were dissected from third instar larvae in M3 Shields and Sang media (Sigma, S3652) supplemented with 5% foetal bovine serum (Sigma, F9665) and 1x Antibiotic-Antimycotic (Gibco, 15240-062). For optogenetic experiments dissections were performed using only a red-light source (LENCENT) for illumination. Salivary glands were subsequently mounted on poly-lysine treated cover slips in dissection media supplemented with 2.5% methyl-cellulose (Sigma, M0387-100G) as in (Gomez-Lamarca et al., 2018).

To image endogenous levels of Notch and CSL in the egg chamber, Notch::YFP flies (Couturier et al., 2012) were crossed with Halo::CSL flies and adult females collected. Dissection of ovaries was performed in in Schneider’s medium (Biowest, L0207-500) supplemented with 15% (v/v) foetal bovine serum (Sigma, F2442), 0.6% (v/v) streptomycin/penicillin antibiotic mix (Gibco, 15140-122), and 0.20 mg/ml insulin (Sigma, I5500). For imaging, tissue was submerged in dissection media on 35-mm poly-D-lysine-coated glass bottom dishes (MatTek, P35GC-1.5-10-C).

Wings were dissected from adult flies three days after eclosion and mounted in glycerol. Images were captured using a Zeis Axioplan 2 microscope with a 5x air objective, Retiga EXi camera, and volocity 6.3.1 software.

## METHOD DETAILS

### Live imaging

Live confocal fluorescence imaging of salivary glands was performed on a Leica SP8 microscope equipped with an Argon laser and 561/633 nm He/Ne lasers using a 63x/1.4 NA HC PL APO CS2 oil immersion objective and two hybrid GaAsP detectors. Nuclei were imaged with an additional 4x zoom. For most experiments, Z-stacks were taken to cover the whole tissue with a step size of 1μm, a resolution of 512x512 px, pinhole set to 3-Airy, 12-bit depth and scanning at 400 Hz. For movies, a single plane was imaged every 2.58 seconds, using the same settings as with Z stacks. We use sequential scanning to first provide excitation wavelengths of 561 nm or 633 nm light for red and far-red fluorescence, and second acquire images of green fluorescent protein using 488 nm excitation or to provide activating blue light for optogenetics (typically 458nm).

Imaging of the ovaries was performed as with the salivary glands with a resolution of 1024x1024 px, pinhole set to 2-Airy, and a Z stack step size of 0.5 μm. Optical zooms for each egg chamber were 5.35, 5.45 and 5.35 respectively (in order from A-C, Fig S3).

### Quantitative RT-qPCR and *in situ* hybridizations

Single molecule fluorescent *in situ* hybridization was performed following the Stellaris protocol as in (Boukhatmi and Bray, 2018). For RT-qPCR, experiments were performed as in Gomez-Lamarca et al (2018). In brief, phenol-chloroform extraction of RNA (using TRI reagent -Invitrogen, AM9738) was performed on 20 salivary gland pairs for each condition and 2 μg of RNA subsequently reverse transcribed into cDNA with 10 U/μL MMLV Reverse Transcriptase, 25 μg/mL oligo(dT)_15_ primers, 1 U/μL RNase inhibitor and 1 mM dNTPs in a 20 μL volume (Promega, M1701 and C110A). Resulting cDNA was then diluted 1:5 to bring Ct values into the measurable range. Quantitative PCR was performed using a Roche Lightcycler 480 II with a 10 μL reaction containing 1 μL cDNA, SYBR green mastermix (Roche) and 0.3 μM primers (Sigma, table 1) targeted against a 142 base pair amplicon in the *E(spl)mβ* coding sequence or control regions (Gomez-Lamarca et al., 2018; Krejčí and Bray, 2007). After amplification samples were denatured, cooled to 65 °C and continuous readings (five per °C) taken as they were reheated to 97 °C to measure the melting point for the primers. Lightcycler software (version 1.5.1) was used to perform a melting curve analysis (to check for non-specific binding) and perform a second derivative maximum analysis to calculate the Ct values for samples. Two technical repeats of the qPCR were completed for each experiment and the mean Ct value taken for analysis, technical repeats with a standard deviation greater than one were excluded. Relative amounts of mRNA (=2^-Ct^)_were then calculated and normalised to the control gene *RpL32* which has previously been found suitable for comparison to the *E(spl)-C* genes (Gomez-Lamarca et al., 2018; Krejčí and Bray, 2007). Replicate numbers refer to the number of biological repeats where the full experimental protocol was completed from start to finish.

**Table 1:**
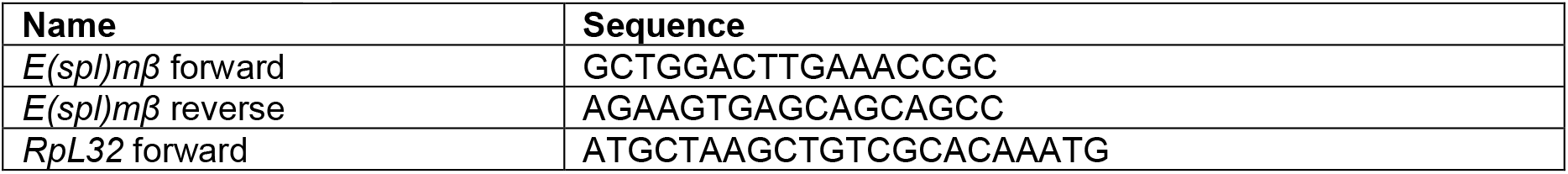
List of Oligonucleotides.

## QUANTIFICATION AND STATISTICAL ANALYSIS

### Image and statistical analysis

Image and video analysis was performed in ImageJ and RStudio (RStudio Team, 2020; Schneider et al., 2012).

For quantifying optogenetic movies and images, regions of interest were manually drawn around the whole cell and nucleus of each cell within a salivary gland. From this the mean nuclear and cytoplasmic fluorescence were calculated. Nuclear accumulation was measured as the mean nuclear fluorescence intensity at each timepoint divided by the initial measurement. The nuclear:cytoplasmic ratio is the mean nuclear fluorescence divided by the cytoplasmic fluorescence.

For quantifying recruitment of CSL to the *E(spl)-C* locus, images were rotated so the locus was a vertical line, a box of width 4μm (45px) and height of the locus tag “band” was drawn and the “plot profile” function of ImageJ used to measure the increased fluorescence across the locus. We aligned all measurements such that the maximum locus tag signal was in the centre of the profile.

